# Updated functional annotation of the *Mycobacterium bovis* AF2122/97 reference genome

**DOI:** 10.1101/757823

**Authors:** Damien Farrell, Joseph Crispell, Stephen V. Gordon

## Abstract

*Mycobacterium bovis* AF2122/97 is the reference strain for the bovine tuberculosis bacillus. We here report an update to the *M. bovis* AF2122/97 genome annotation to reflect 616 new protein identifications which replace many of the old hypothetical coding sequences and proteins of unknown function in the genome. These changes integrate information from functional assignments of orthologous coding sequences in the *Mycobacterium tuberculosis* H37Rv genome. We have also added 69 additional new gene names.

## Background

*Mycobacterium bovis (M. bovis)* is a causative agent of bovine tuberculosis (bTB) and the most widely studied animal-adapted member of the *Mycobacterium tuberculosis* complex (MTBC). The genome of the *M. bovis* AF2122/97 strain was first sequenced in 2003 [1] and is considered to be the reference sequence for this species. *Mycobacterium bovis* has considerable importance as the basis for comparative studies into animal- and human-adapted species of the MTBC. This genome was revised in 2017 [2] with an updated sequence that included the previously missing RD900 region and added 42 new coding sequences. These revisions brought the annotation in line with updates to the *M. tuberculosis* H37Rv genome, to which *M. bovis* shares high identity.

### Hypothetical and unknown proteins

Proteins encoded by genes with no clear functional activity have been traditionally annotated with designations such as ‘unknown protein’, ‘hypothetical protein’ or ‘conserved hypothetical’. These labels are usually placed in the /product field of the annotation file (in this context we refer to the genbank file format). This /product qualifier is not automatically updated when new protein functions are discovered, and hence genome annotation files become out of date over time if not regularly updated. Cross references to databases with functional information are automatically added in a /db_xref qualifier during updates on the DDBJ/EMBL/GenBank system; however these are not human readable. Therefore, it is a valuable and necessary exercise to update the protein product information directly in genome annotations of reference species.

### New sources of annotation

Since the original annotation of *M. bovis* AF2122/97, the function of many hypothetical and newly identified proteins has been recognised. These data were collated by Doerks at al. in 2012 [3] who added approximately 620 new functional assignments to the remaining unknowns in the *M. tuberculosis* H37Rv reference genome. This was done using orthology and genomic context evidence. If a hypothetical protein was a member of a known orthologous group in the eggNOG database, this annotation was transferred to the corresponding *M. tuberculosis* protein. The STRING tool was also used, combining gene fusion events and significant co-occurrence to predict links with known proteins. The annotations vary in specificity; those predicted through association are more ‘functional hints’ as to the nature of the protein function rather than ascribing a specific function.

The PATRIC database [4] have used their own pipeline based on RASTtk to re-annotate H37Rv. The PATRIC annotation has some additional functional assignments and small genes that were not in the *M. tuberculosis* H37Rv reference. Some of the genes found by Doerks et al. are also assigned in PATRIC and a few have since been assigned to the reference. Figure 1 shows the overlap in these two sets with the existing unknown proteins in the H37Rv reference.

**Figure 1:**
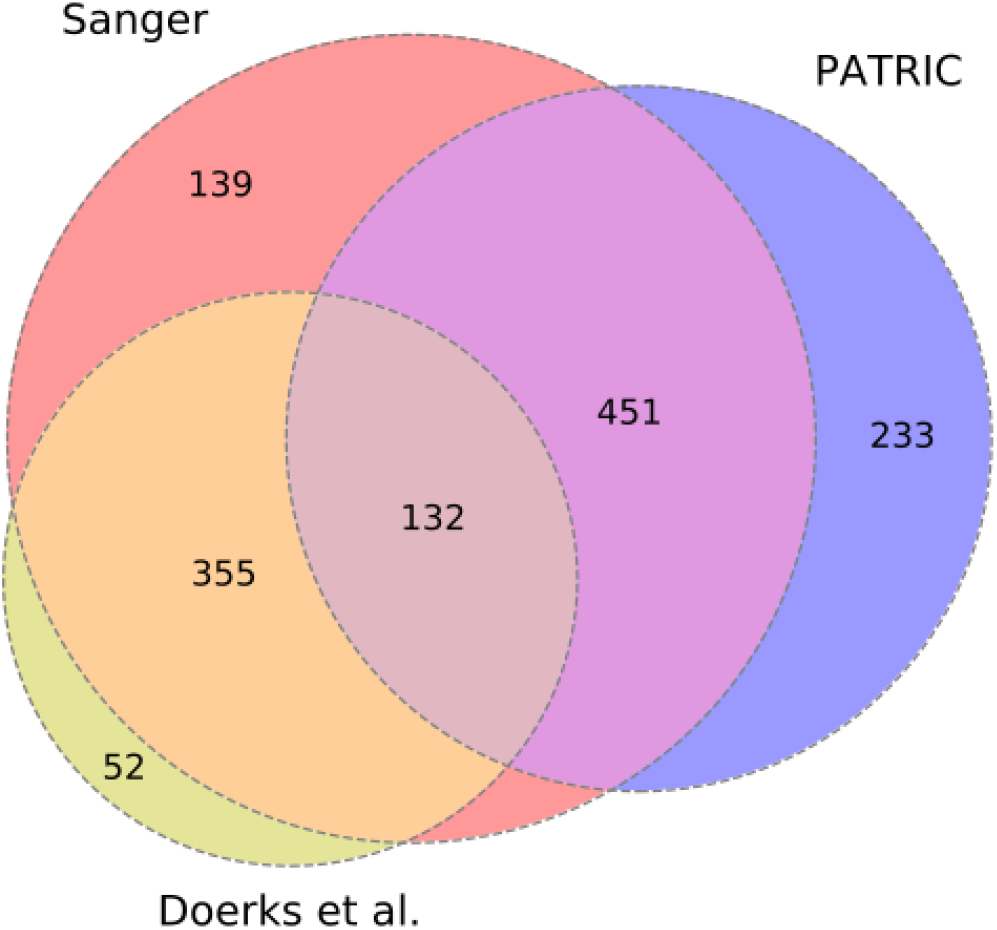
Overlap between the hypothetical/unknown proteins in the reference (Sanger) and PATRIC H37Rv annotations and those found by Doerks et al.

UniProt [5] stores up-to-date protein annotations for the *M. tuberculosis* H37Rv strain, several of which are recent and were not present in the either PATRIC or Doerks at al. data. These three sources were used to make an integrated table that was used to update the protein product field of the *M. bovis* AF2122/97 genome annotation.

## Methods

Data from three sources was used: 1) Doerks et al. (2012); 2) the PATRIC H37Rv annotation (https://www.patricbrc.org/view/Genome/83332.12); and 3) the UniProt *M. tuberculosis* H37Rv proteome (https://www.uniprot.org/proteomes/UP000001584) were downloaded as csv files and combined together by matching corresponding entries on the H37Rv locus tags (/locus_tag field). For all the unknown proteins in the H37Rv genome we selected updated annotations from these combined sources: if an annotation was present in the Doerk dataset this was used preferentially, then PATRIC and finally UniProt. The order of preference was not significant. The majority of the data was provided from the Doerks dataset though 62 proteins from this dataset were excluded due to lack of a specific functional description.

The resulting table with new protein products was then matched to the *M. bovis* orthologous genes using a mapping between *M. tuberculosis* H37Rv and *M. bovis* locus tags. In a final step we performed a BLAST [6] of the remaining unknowns to the Protein Data Bank and found five additional proteins that have structures and function ascribed which were also added. Analysis was done in Python, utilising the Biopython [7] and pandas [8] libraries.

## Results

The current *M. bovis* genome annotation contains 3989 protein coding genes [2]. Of these 1097 were marked as hypothetical, conserved or unknown proteins. These data are summarized in Table 1, with two *M. tuberculosis* H37Rv annotations for comparison. We have now added a total of 616 new protein product annotations to the genome which includes five products from a PDB search. 69 new gene names were added from the UniProt data. There are now 488 hypothetical/unknowns remaining in the updated *M. bovis* AF2122/97 annotation. This revised annotation has been submitted to DDBJ/ENA/GenBank and is available under the accession no. <u>LT708304</u>.

**Table 1:** Current annotation statistics for the *M. bovis* and H37Rv genomes. The revised numbers for the updated annotations for AF2122/97 are shown in brackets.

|  | Coding sequences | CDS with gene names | Hypothetical | Pseudogenes |
| --- | --- | --- | --- | --- |
| **Mbovis AF2122/97** | 3989 | 1964 (2026) | 1097 (488) | 11 |
| **MTB-H37Rv (Reference)** | 4018 | 1953 | 1097 | 27 |
| **MTB-H37Rv (Broad)** | 4143 | 16 | 843 | 92 |

## Conclusion

Reference genomes stored on the INSDC (International Nucleotide Sequence Database Collaboration) provide the primary sources of annotation for virtually all bacterial species. Though there are now multiple alternative information sources for bacterial genomes, most are specialist databases and not universally known. The proliferation of data sources risks fragmentation of genome annotation and linked functional information; it is vitally important that reference sequence annotation in GenBank and Ensembl, the first port of call for the majority of researchers, are as up to date as possible. The *M. bovis* AF2122/97 strain is well established as a reference for *M. bovis* and the MTBC, and will remain a research cornerstone for the foreseeable future; maintaining an updated genome annotation to the research community drove our current work. We note that the latest reference annotation of *M. tuberculosis* H37Rv also lacks many of the updates we have added here to hypothetical proteins, underlining the need for constant curation of reference sequence annotation.

## Data Availability

All data sources, output files and the Jupyter notebook used to produce this analysis are stored in a github repository at the following url: https://github.com/dmnfarrell/gordon-group/tree/master/mbovis_annotation.

## Acknowledgements

This work was supported by the Irish Department of Agriculture Food and the Marine grant 15/S/651 (NEXUSMAP) and Science Foundation Ireland (SFI) grant 16/BBSRC/3390, part of the SFI–Biotechnology and Biological Sciences Research Council joint funding partnership.

## Notes

https://github.com/dmnfarrell/gordon-group/tree/master/mbovis_annotation

## References

[1] T. Garnier et al., “The complete genome sequence of Mycobacterium bovis.,” Proc. Natl. Acad. Sci. U. S. A., vol. 100, no. 13, pp. 7877–82, Jun. 2003.

[2] K. M. Malone, D. Farrell, T. P. Stuber, and O. T. Schubert, “Updated Reference Genome Sequence and Annotation of Mycobacterium bovis AF2122/97,” Am. Soc. Microbiol., vol. 5, no. 14, pp. 17–18, 2017.

[3] T. Doerks, V. van Noort, P. Minguez, and P. Bork, “Annotation of the M. tuberculosis hypothetical orfeome: adding functional information to more than half of the uncharacterized proteins.,” PLoS One, vol. 7, no. 4, p. e34302, Jan. 2012.

[4] A. R. Wattam et al., “PATRIC, the bacterial bioinformatics database and analysis resource,” Nucleic Acids Res., 2014.

[5] A. Bateman et al., “UniProt: The universal protein knowledgebase,” Nucleic Acids Res., 2017.

[6] S. F. Altschul, W. Gish, W. Miller, E. W. Myers, and D. J. Lipman, “Basic local alignment search tool,” J. Mol. Biol., no. 215, pp. 403–410, 1990.

[7] P. J. A. Cock et al., “Biopython: Freely available Python tools for computational molecular biology and bioinformatics,” Bioinformatics, 2009.

[8] W. Mckinney, “Pandas, Python Data Analysis Library,” 2015. [Online]. Available: http://pandas.pydata.org/.

